# CRISPR editing of candidate host factors that impact influenza A virus infection

**DOI:** 10.1101/2024.09.10.612185

**Authors:** Pyae Phyo Kyawe, Ping Liu, Zhaozhao Jiang, Evan S. Bradley, Thomas Cicuto, Melanie I. Trombly, Neal Silverman, Katherine A. Fitzgerald, William M. McDougall, Jennifer P. Wang

**Affiliations:** Diabetes Center of Excellence, Department of Medicine, University of Massachusetts Chan Medical School; Interdisciplinary Graduate Program, Morningside Graduate School of Biomedical Sciences; Division of Innate Immunity, Department of Medicine, University of Massachusetts Chan Medical School; Department of Microbiology, University of Massachusetts Chan Medical School; Division of Infectious Diseases and Immunology, Department of Medicine, University of Massachusetts Chan Medical School

**Author notes:** **Byline** Influenza A virus, sialic acid, host factors, ADAR, interferon. **Correspondent footnote:** Corresponding author: Jennifer P. Wang, MD, Department of Medicine, Diabetes Center of Excellence, University of Massachusetts Chan Medical School, ASC 7-2047, 368 Plantation St., Worcester, MA 01605.

## Abstract

Influenza A virus (IAV) is a respiratory pathogen with a segmented negative-sense RNA genome that can cause epidemics and pandemics. The host factors required for the complete IAV infectious cycle have not been fully identified. Here, we examined select host factors that were identified by independent CRISPR screens as candidate contributors to IAV infectivity. We performed CRISPR-mediated knockout of cytidine monophosphate N-acetylneuraminic acid synthetase (CMAS) as well as CRISPR-mediated overexpression of beta-1,4 N-acetylgalactosaminyltransferase 2 (B4GALNT2) and adenosine deaminase acting on RNA 1 (ADAR1) in the human bronchial epithelial A549 cell line and evaluated IAV infectivity. We confirmed that the knockout of *CMAS* or overexpression of *B4GALNT2* restricts IAV infection by diminishing binding to the cell surface but has no effect on vesicular stomatitis virus infection. While ADAR1 overexpression does not significantly inhibit IAV replication, it has a pro-viral effect with coxsackie B virus (CVB) infection. This pro-viral effect is not likely secondary to reduced type I interferon (IFN) production, as the induction of the IFN-stimulated genes *ISG15* and *CXCL10* is negligible in both parent and ADAR1-overexpressing A549 cells following CVB challenge. In contrast, *ISG15* and *CXCL10* production is robust and equal for parent and ADAR1-overexpressing A549 cells challenged with IAV. Taken together, these data provide insight into how host factors identified in CRISPR screens can be further explored to understand the dynamics of pro- and anti-viral factors.

**Importance:** Influenza A virus (IAV) remains a global threat due to its ability to cause pandemics, making the identification of host factors essential for developing new antiviral strategies. In this study, we utilized CRISPR-based techniques to investigate host factors identified in screens as reducing IAV infectivity. Knockout of CMAS, a key enzyme in sialic acid biosynthesis, significantly reduced IAV binding and infection by disrupting sialic acid production on the cell surface. Overexpression of B4GALNT2 had similar effects, conferring resistance to IAV infection through diminished cell-surface binding. While overexpression of ADAR1, known for its role in RNA editing and immune regulation, slightly reduced IAV replication, it increased coxsackie B virus replication. Such findings reveal the diverse roles of host factors in viral infection, offering insights for targeted therapeutic development against IAV and other pathogens.

## Introduction

Influenza A virus (IAV) is an enveloped virus with a segmented negative-sense single-stranded RNA genome that is a recurring respiratory pathogen in humans. Understanding how IAV infects cells and identifying the host components necessary for productive infection is essential given its tremendous potential for global illness, such as with pandemic outbreaks (1). The 2009 influenza pandemic and the recent SARS-CoV-2 pandemics showed how the spread of respiratory pathogens is driven by socioeconomic issues and can overwhelm the healthcare system; therefore, a finer understanding of IAV-host dynamics is warranted.

Various RNA-based genome-wide screening methods, mainly arrayed small interfering RNA (siRNA) and pooled short hairpin RNA (shRNA)-based screens, have been used to identify essential host factors necessary for productive IAV infection (2-7). In lentiviral-based pooled shRNA screens, the time required to achieve sufficient knockdown of the target genes is usually longer compared to arrayed siRNA-based screens. Furthermore, the requirement to carry cells for longer also rules out host factors usually involved in cell survival or proliferation. Among candidates detected in the screens, fewer than 10% were shared across screens. This lack of overlap could be due to the differences in virus strains, source of reagents, knockdown efficiency, host cell lines, endpoints, and methods of detection (reviewed in (8)) and underscores the potential for false positives with these RNAi approaches.

CRISPR-Cas9 systems minimize the drawbacks of using RNAi-based systems as they have fewer off-target effects and allow for the generation of permanent homozygous null lines (9-11). CRISPR-Cas9-based screening methods have become a standard format for genome-wide screens for host factors during virus infections due to ease of scalability and minimal off-target effects. Many CRISPR-based genome-wide knockout (KO) screens have been performed for viruses, including IAV (12-14), coronaviruses (15-17), flaviviruses (18-20), and HIV (21-23). Han et al. first utilized the CRISPR-Cas9-based gene KO screening strategy in A549 cells to determine host factors essential for infection of avian influenza virus (13). Next-generation sequencing of survivor cells identified an enrichment of host factors involved in sialic acid biosynthesis (including *CMAS* encoding cytidine monophosphate N-acetylneuraminic acid synthetase), sialic acid transport (*SLC35A1, SCL35A2* encoding members of solute carrier family 35), and glycan modification/processing (including *B4GALNT4* encoding beta-1,4-N-acetyl-galactosaminyltransferase 4) (13). Using a CRISPR-Cas9-based screen, Yi et al. also identified several sialic acid-related genes, including *SLC35A1, GNE, CMAS,* and *NANS*, as critical for IAV infection (24). However, a downside of CRISPR-Cas9’s ability to generate a homozygous null phenotype also means that genes required for viability and proliferation are eliminated from the screen, since cells with such KO genes would not have survived or expanded. Candidates identified are those that confer strong resistance to virus infection, but survivor screens are not designed to detect candidates that confer moderate resistance to virus infection. Furthermore, pro-viral host factors will not be detected in a survivor screen.

The development of catalytically inactive, endonuclease-deficient Cas9 (dCas9) fused to transcriptional repressor domains, such as KRAB, permits further study of gene function without generating a potentially cell lethal phenotype. Furthermore, manipulating genes at the transcriptional level rather than generating indels on the genomic DNA results in fewer off-target effects, thereby decreasing the chances of false positives or negatives (25, 26). dCas9 can also be fused to transcriptional activators (e.g., VPR, VP64, p65, HSF) to induce overexpression through guide RNA (gRNA)-directed transcriptional expression instead of conventional cDNA-mediated overexpression (27, 28). Recently, a CRISPR-activation (CRISPRa) screen performed to identify host factors that inhibit IAV infection when overexpressed showed that overexpression of *B4GALNT2* (encoding beta-1,4-N-acetyl-galactosaminyltransferase *2*, a participant in sialic acid modification) results in inhibitory activity against influenza viruses with α2,3-linked sialic acid receptor preference (29). Using recombinant IAV harboring single gRNA (sgRNAs) to infect A549 cells expressing dCas9-VP64 and MS2-p65-HSF1, which allowed for enrichment of recombinant viruses carrying pro-viral sgRNAs beneficial for multiple rounds of infection, King et al. identified *TREX1,* which encodes three prime repair exonuclease, as a pro-viral factor that degrades mitochondrial DNA released into the cytoplasm during IAV infection, preventing the activation of DNA-sensing pathways and diminishing type I interferon (IFN) (30).

To identify host factors that confer moderate resistance to virus infection as well as pro-viral host factors, we generated A549 cells expressing a human CRISPR interference (CRISPRi) Dolcetto library and a human CRISPR activation (CRISPRa) Calabrese library and performed whole-genome arrayed CRISPRa/i screens to identify host protein-coding genes that diminish IAV infection. While validation of most candidates is being pursued in a more detailed report to be published separately, here, we validated the role of one CRISPRi candidate, *CMAS*, in IAV infection. We also demonstrated how Cas9-driven overexpression of *B4GALNT2* using CRISPRa reduced IAV infection through impaired sialic acid production. Finally, we examined the impact of overexpressing the CRISPRa candidate *ADAR1* (adenosine deaminase acting on RNA 1) in the context of IAV and other virus infections.

## Materials and Methods

### Viruses

IAV strain WSN/33 (H1N1) was obtained from A. L. Brass (3), strain Puerto Rico/8/1934 (PR/8) and Hong Kong/1968 (HK/68) from Charles River Laboratories, and California/04/2009 (pdm2009) from American Type Culture Collection (ATCC, #VR-1805). VSV strain Indiana was obtained from ATCC (#VR-1238). CVB4 strain JVB was obtained from ATCC (#VR-184).

### Cell lines

The following cell lines were obtained from ATCC: A549 (#CCL-185), Madin-Darby canine kidney (MDCK) cells (#CCL-34), HeLa (#CCL-2), and baby hamster kidney (BHK-21) cells (#CCL-10). Cells were grown in Dulbecco’s Modified Eagle Medium (DMEM) supplemented with 10% fetal bovine serum, 100 U/mL penicillin, 100 μg/mL streptomycin, and GlutaMax^TM^. The cell lines were routinely monitored for mycoplasma. HEK293T and HEK293T ADAR1 KO cells were a gift from Dr. Daniel Stetson (31).

A549 Cas9 and A549-dCas9-VP64 cells were generated via lentiviral transduction of SpCas9 or the VP64 activator domain (dCas9-VP64) into A549 cells. Clonal cell lines with high expression of Cas9 or dCas9-VP64 proteins were established by limiting dilution after selection with 10 μg/mL blasticidin S HCl (ThermoFisher #A1113903) for 14 d. A549 CMAS-deficient cell lines were generated by transfecting A549-Cas9 cells (32) with pGS-CMAS-Neo plasmid with sgRNA targeting exon 4 of CMAS with CCATCCCAGTCTTGTCGACG gRNA from a U6 promoter and a neomycin-resistance marker (GenScript). Clonal A549 CMAS-deficient cell lines were established by limiting dilution after selection with 800 μg/mL G418 (Geneticin^TM^, ThermoFisher #10131027) for 14 d.

For A549 Cal^B4GALNT2^ and Cal^ADAR1^ overexpressing lines (cell lines were named “Cal” after the Calabrese library with the overexpressed gene in superscript), sgRNA sequences for the CRISPRa candidate genes were cloned into expression vector pXPR-502 as described previously (28, 33). sgRNA targeting the B4GALNT2 promoter region or ADAR1 and complementary oligos with appropriate nucleotide overhangs (sgRNA for B4GALNT2: GGGCAAATTCTCGGCGAGTA, ADAR1 g1: AACCGGCCTGAAACCAAGCG, g2: CTTCCGTAGTTCTCATGCAG, g3: CGCTGCATGAGAACTACGGA) were ordered from ThermoFisher and annealed with a thermocycler to form dsDNA duplexes. Specifically, oligos were mixed with T4 Ligase Buffer (NEB) and T4 PNK enzyme and annealing was performed with the thermocycler following the program: 37 °C for 30 min, 95 °C for 5 min, then decreasing the temperature by 5 °C/min until 25 °C was reached. 90 μL of water was added to each reaction and 2 μL of duplexed oligos were ligated to 25 ng of linearized pXPR-502 lentiviral expression vector in 20 μL ligation reaction at 16 °C overnight. 7 μL of the ligated material was used to transform Stbl3 *E. coli*, which was plated on LB plates containing 100 μg/mL ampicillin. At least three bacterial colonies were sent out for Sanger sequencing and plasmids with the correct sgRNA sequences were expanded for lentiviral transduction (33). Confluent HEK293T cells in a 10 cm dish were transfected with plasmids expressing VSV-G envelope protein (pMD2.G), HIV structural proteins (psPAX2), and pXPR-502 expressing the specific sgRNA from a U6 promoter and a puromycin resistance gene from an EF-1a promoter at the ratio of 1:1.5:3 (4, 6, and 12 μg, respectively) using TransIt-293T. Supernatants were collected every 24 h for 3 d and pooled. The supernatant was concentrated with LentiX-concentrator and added to A549-dCas9-VP64 cells. Lentivirus-transformed cells were then selected with 2 μg/mL puromycin for 7 d. The survivor cells were then expanded and maintained in complete DMEM supplemented with 2 μg/mL puromycin. Supernatants were collected for 3 d and passed through a 0.45 μm filter to remove debris before being added to target cell lines. Media was replaced with DMEM + 10 μg/mL blasticidin, 2 μg/mL puromycin (A1113803, Invitrogen) at 24 h post-transduction and cells were cultured for 7 d. Clonal knockdown and overexpression cell lines were established by limiting dilution and with passage in selective media.

### Western blots

Cells were lysed with RIPA buffer with protease inhibitor cocktail (Sigma #P8340) for 10 min at room temperature. The cell lysate was centrifuged at 20,000 x g for 10 min to remove cell debris. The concentration of the lysate was determined by BCA assay (ThermoFisher #PI23227). 35 μg of protein lysate was heated at 95 °C for 5 min and loaded onto Novex Tris-Glycine Mini-Protein Gel 4–12% (ThermoFisher #XP04200BOX) and run at 100V for 1.5 h. The proteins were then transferred onto a PVDF membrane (ThermoFisher #88518) with an Xcell II Blot Module (ThermoFisher #EI0002) for 1 h at 20V. The membrane was blocked with 5% nonfat dry milk in TBS+0.1% Tween 20 (TBS-T) for at least 1 h. The membrane was incubated with primary antibody – mouse anti-CMAS (Millipore Sigma #SAB1405174) diluted 1:1000 in 5% nonfat dry milk in TBS-T or rabbit anti-ADAR (Cell Signaling #81284S) diluted 1:1000 in 5% nonfat dry milk/TBS-T – overnight with constant rocking. Membranes were washed three times with TBS-T and incubated with HRP-conjugated secondary antibody – horse anti-mouse IgG-HRP (Cell Signaling #7076S) diluted 1:1000 in 5% nonfat dry milk/TBS-T or goat anti-rabbit IgG-HRP (Vector Laboratories #PI-1000) diluted 1:20,000 in 5% nonfat dry milk/TBS-T – for 1 h at room temperature. Membranes were washed 5 times with TBS-T and then visualized with SuperSignal West Pico PLUS Chemiluminescent Substrate (ThermoFisher #34580) using a BioRad Imager. Membranes were stripped with Restore Plus Western stripping buffer (ThermoFisher #46430) for 10 min and washed twice with TBS-T. The membrane was then blocked in 3% BSA in TBS-T for 1 h and then incubated with anti-actin HRP (Santa Cruz Biotechnology #sc-37778) diluted 1:2000 in 3% BSA for 1 h. The membrane was washed five times in TBS-T and visualized.

### RT-qPCR

RNA was extracted from cell lines with TRIzol and 1 μg of RNA was converted to cDNA using the Qiagen reverse transcription kit. Equal concentrations of cDNA were loaded for quantification of target gene levels by SYBR Green qPCR using HPRT as a housekeeping gene for 40 cycles. Fold change was calculated based on the 2^ΔΔCT^ method. Primers are as follows (Life Technologies Corporation, ThermoFisher):

*CMAS* F: ACAAGACTGGGATGGAGAA, R: ACTATGTTCAGCTCGCATTT; *B4GALNT2* F: CTACGATGGAATCTGGCTGTT, R: GCCATAGGCATCCTGAAAGT; *ADAR1* F: ATCAGCGGGCTGTTAGAATATG, R: AAACTCTCGGCCATTGATGAC; *HPRT* F: ATCAGACTGAAGAGCTATTGTAATGA, R: TGGCTTATATCCAACACTTCGTG; *ISG15* F: CGCAGATCACCCAGAAGATCG, R: GATCTCAGAAATACCCCAGCCA; and *CXCL10* F: TTCTGATTTGCTGCCTTATCTTTC, R: TTCTTGATGGCCTTCGATTCTGG.

### RNA-sequencing and analysis

RNA was purified from cell lines by TRIzol Reagent. Strand-specific total RNA with ribosomal RNA depletion (1 µg of input RNA) was used to generate libraries generated with the TruSeq Stranded Total RNA Sample Prep Kit (Illumina) and sequenced on an Illumina NextSeq 2000 machine. Paired-end sequence reads were aligned to the human reference genome (hg38_v44) using STAR (34) and transcript quantification was performed with RSEM (35) through the UMass Chan Medical School DolphinNext (36). Downstream analysis was performed in R version 4.4.0 using the RStudio interface. Relative transcript abundance was estimated using DESeq2 based on normalized counts of alignments, three replicates of parental A549-dCas9-VP64 were compared to clones overexpressing either ADAR1 or B4GALNT2 (37). The cutoff for significance was p < 0.1 adjusted for multiple comparisons. Volcano plots were constructed using the EnhancedVolcano package (38).

### Lectin staining and flow cytometry

Cells were trypsinized at 37 °C for 5 min to obtain single-cell suspensions and transferred to round bottom 96-well plates at 2 x 10^5^ cells per well. Cells were washed with 200 μL of PBS twice. Biotinylated lectins *Maackia amurensis* (MAA, 2 μg/mL, Vector Laboratories #B-1265-1), *Sambucus nigra* (SNA, 2 ug/mL, Vector Laboratories #B-1305-2), or *Dolichos biflorus* agglutinin DBA (2 μg/mL, Vector Laboratories #B-1035-5) were added to each well in 100 μL volume and binding was performed for 1 h at 4 °C. Cells were washed twice with PBS to remove the unbound lectins and Alexa Fluor 647-conjugated streptavidin (SA-647, ThermoFisher Scientific #S21374) was added to each well at a working concentration of 2 μg/mL for 1 h at 4 °C. Cells were washed twice with PBS and fixed with 4% paraformaldehyde (PFA) for 10 min at 4 °C and then washed twice with PBS and resuspended in 150 μL PBS for flow cytometry. At least 10,000 events were captured and analyzed on a BD FACSCelesta running BD FACS Diva Software with a standard laser and filter combination. All data were visualized and analyzed with FlowJo software.

### IAV infection and TCID_50_ assay

Cells were plated at 1.5 x 10^4^ cells/well in 100 μL media in 96-well black plates with a clear bottom (Corning #3603) and incubated overnight at 37 °C. The cells were then challenged with virus diluted in 100 μL serum-free DMEM for 1 h at 37 °C. Cells were washed to remove unbound virus, and 100 μL of fresh growth media was added to each well. Plates were incubated for 24 h at 37 °C and supernatants were collected and saved at -80 °C for 50% tissue culture infectious dose (TCID_50_) assays. Cells were fixed in 4% PFA for 15 min at room temperature and permeabilized with 0.2% Triton X-100 in PBS for 15 min at room temperature. The cells were then stained with anti-HA antibody at 1:20 (hybridoma HA36-4-5.2, Wistar Institute), followed by Alexa Fluor 488 goat anti-mouse secondary antibody at 1:1000 (A11001, Invitrogen) or Alexa Fluor 647 goat anti-mouse secondary antibody at 1:1000 (A-21235, Invitrogen). For some experiments, anti-NP antibody (Millipore Sigma, MAB8258, 1:1000 in PBS+1% BSA) followed by goat anti-mouse IgG-conjugated to Alexa Fluor 647 (ThermoFisher #A21235 1:1000 in PBS+1% BSA) was used. Cells were imaged using the Celigo Imaging Cytometer (Nexcelom) and HA- or NP-positive cells normalized to the total number of cells (determined by DAPI, Invitrogen #D1306) were analyzed using the “Expression Analysis” on the Celigo Imaging Cytometer.

For multi-cycle viral replication assays, cells were seeded in 24 well plates, infected, and supernatants were collected at 1, 24, and 48 h post-infection (hpi) and stored at -80 °C. Samples were titered on MDCK cells using a standard TCID_50_ assay as described elsewhere (39). In brief, MDCK cells were plated at 2 x 10^5^ cells/well in 96-well plates in 100 μL growth media and incubated overnight at 37 °C. Supernatants were thawed on ice and ten-fold serial dilutions of viruses were prepared in serum-free DMEM. MDCK cells were washed twice with PBS to remove FBS, and then 100 μL of diluted viruses were added to MDCK cells in replicates of eight. The cells were then incubated at 37 °C for 1 h with gentle shaking. Virus dilutions were removed by aspiration, and the cell monolayers were then washed twice with PBS to remove unbound virus. Cells were then maintained in DMEM supplemented by 0.2% BSA + 0.5 μg/mL tosyl phenylalanyl chloromethyl ketone (TPCK)-treated trypsin (Sigma #T8802) and monitored for cytopathic effect daily. After 72 h, the TCID_50_/mL was calculated using the Reed-Muench method. For some experiments, virus was titered by plaque assay using MDCK cells as previously reported (40).

### Sialidase treatment and IAV infection of A549 cells

A549 cells (2 x 10^4^ cells/well) were seeded in 96-well plates and incubated overnight at 37 °C. The cells were treated with various concentrations of *C. perfringens* sialidase (Sigma Aldrich #11585886001) for 1 h at 37 °C, washed, then infected with IAV WSN/33 (MOI 50) for 8 h. The percentage of NP+ cells was determined by NP immunofluorescence normalized to total cell count (nuclear staining).

### VSV infection and plaque assay

Cells were plated and infected with VSV, then supernatants were collected 24 hpi and titered on BHK-21 cells as described previously (41). Ten-fold serial dilutions of supernatants in serum-free DMEM were added to confluent BHK-21 cells in 12-well plates in a final volume of 100 μL/well in duplicates. Cells were then incubated at 37 °C for 1 h with gentle shaking. Supernatants were removed by aspiration, and the cell monolayers were washed with serum-free DMEM. A 2% agarose overlay with 1X Modified Eagle’s Medium (Gibco #11935) was added to each well and allowed to solidify at room temperature and then incubated at 37 °C for 24 h. Cells were fixed/stained for at least 2 h at room temperature with 0.5% crystal violet dissolved in 4% PFA. Overlays were removed, and the PFU/mL was calculated by averaging the number of plaques in the duplicate wells and dividing by the product of the dilution factor and volume added to each well.

### CVB infection and plaque assay and poly I:C treatment

Cells were plated and infected with CVB4, and then supernatants were collected 24 hpi and titered on HeLa cells based on standard protocols. Ten-fold serial dilutions of supernatants in serum-free DMEM were added to confluent HeLa cells in 6-well plates in a final volume of 100 μL/well in duplicates. Cells were then incubated at 37 °C for 1 h with gentle shaking. Supernatants were removed by aspiration and the cell monolayers were washed with serum-free DMEM. An overlay mixture (1% agar, 1X Modified Eagle’s Medium, 5% FBS, and 5 mM MgCl_2_) was added to each well and allowed to solidify at room temperature and then incubated at 37 °C for 48 h. 500 μL of 2X MTT/INT dye (thiazolyl blue tetrazolium bromide/iodonitrotetrazolium chloride) was added on top of each overlay, and the plates were incubated at 37 °C for at least 2 h. PFU/mL was calculated by averaging the number of plaques in the duplicate wells and dividing by the product of the dilution factor and volume added to each well. High-molecular-weight (HMW) poly I:C (Invivogen #tlrl-pic) was added to cells for 24 h.

### Fluorescence *in situ* hybridization

Cells were seeded onto rat tail collagen-coated coverslips in 12-well plates at 10,000 cells/well in 1 mL complete DMEM overnight. Cells were then washed twice with serum-free DMEM and prechilled on ice for 45 min. IAV WSN/33 diluted in DMEM/0.2% BSA was added to each well at an MOI of 1 and binding was performed on ice for 1 h. Unbound virus was washed off with serum-free DMEM and pre-warmed DMEM+FBS was added to each well to facilitate endocytosis and cells were kept at 37 °C for 20 min. Cells were fixed with 4% PFA for 10 min at room temperature and permeabilized with 70% ethanol for 2 h at 4 °C. Hybridization was performed using Quasar 570-conjugated fluorescence *in situ* hybridization (FISH) probes against segment 7 of WSN/33 viral RNA (vRNA, sequences available at (42)) following the manufacturer’s protocol (BioSearch Technologies). Briefly, cells were washed with Wash Buffer A for 5 min, and then a fluorescence in situ hybridization (FISH) probe in hybridization buffer was added for 12 h at 37 °C. The probes were removed by washing the cells with Wash Buffer A for 30 min at 37 °C in the dark. DAPI (5 ng/mL diluted in Wash Buffer A) was added to each cell and stained in the dark for 30 min at 37 °C. DAPI was removed, and cells were incubated in Wash Buffer B for 5 min at room temperature. Coverslips were mounted with ProLong^TM^ Diamond Antifade Mountant (ThermoFisher #P36970). ImageJ was used to determine the average mean fluorescence intensity (representing the average intensity of WSN/33 segment 7 vRNA staining) in each cell.

### IAV-binding assay

Trypsinized cells were added to 96-well round bottom plates at 2 x 10^5^ cells per well and chilled on ice for 1 h at 4 °C. Cells were washed with cold PBS and then IAV WSN/33 was added to the prechilled cells at the indicated MOI in 100 μL serum-free DMEM. IAV binding was performed on ice for 1 h. Cells were washed twice with cold PBS to remove unbound virus, then fixed with 4% PFA for 10 min at 4 °C. Cells were then washed with PBS twice before staining with anti-HA antibody (hybridoma HA36-4-5.2, Wistar Institute, 1:100 dilution in PBS/1% BSA) for 1 h at 4 °C. Cells were then washed with PBS twice and stained with Alexa Fluor 647-conjugated goat anti-mouse antibody (ThermoFisher #A21235) diluted 1:400 in PBS/1% BSA for 1 h at 4 °C. Cells were then washed twice with PBS and resuspended in 150 μL PBS for flow cytometry.

### Microscopy

Samples were imaged by confocal microscopy using a Leica SP-8 Stellaris System.

### Statistical analysis

Statistical analyses were performed using GraphPad Prism Version 10.1.1. Data are presented as mean ± standard deviation. Statistical tests were used as indicated in the figure legends. p < 0.05 was considered statistically significant.

## Results

### CRISPR interference and activation screening strategy

For an arrayed screening strategy, summarized in **Figure S1,** clonal A549-dCas9-VP64 cells (for CRISPRa screening) and A549-dCas9-KRAB cells (for CRISPRi screening) were selected based on cell growth properties and high Cas9 expression. For the CRISPRa screen, the Calabrese whole-genome library was introduced into the A549-dCas9-VP64 cell line. For the CRISPRi screen, the Dolcetto whole-genome library was introduced into the A549-dCas9-KRAB cell line. Next-generation sequencing data to assess the reads per gRNA demonstrated that the gRNA libraries were represented in each system at >95%. Several optimization steps were performed prior to screening, including establishing the optimal conditioned media concentration for fluorescence-activated cell sorting for single cells into 384-well plates. IAV strain WSN/33 was used at an MOI of 0.3, yielding ∼40–50% infectivity at 24 hpi. For each screen, 1000 plates were infected with IAV WSN/33. Wells containing at least 5000 cells and with evidence of decreased infection by HA immunostaining at 24 hpi were selected for genomic DNA isolation and sequencing of amplicons containing gRNAs. gRNA matches were identified in 75% of candidate wells. Genomic DNA was isolated and sequenced with the top ten candidates shown in **Table S1**.

Two genes involved in *de novo* sialic acid biosynthesis, *CMAS*, and *SLC35A1*, were identified as reducing IAV WSN/33 infection in our CRISPRi screen, with *CMAS* appearing twice (**Table S1** and (43)). CMAS converts N-acetylneuraminic acid to cytidine-5-monophosphate (CMP)-sialic acid in the nucleus, which is subsequently transported from the cytosol to the Golgi by the Golgi-resident membrane transporter SLC35A1 (44, 45). Supporting these findings, both *CMAS* and *SLC35A1* were separately identified in independent screens (13, 46). Other groups have identified CMAS as important for attachment and entry of swine, avian, and human IAVs (47). ADAR1, an RNA-binding ISG, was among the top hits of the CRISPRa screen (see (43) and **Table S1**). ADAR1 is a dsRNA-binding protein that converts adenosine to inosine (referred to as A-to-I editing) in endogenous dsRNA to prevent aberrant IFN activation. ADAR1 has been proposed to have both pro- and anti-viral functions depending on the cell lines and viruses tested (48). There are two isoforms of ADAR1, the IFN-inducible cytoplasmic p150 and the constitutive nuclear p110. The sgRNA sequenced in the screen was directed to the ADAR1 p150 promoter. Validation of the impact of *CMAS* deficiency and *ADAR1* overexpression on IAV infection is pursued below.

### Targeting of CMAS by CRISPRi in A549 cells inhibits infection with IAV but not VSV

To confirm the role of CMAS during IAV infection, we generated CMAS KO cells by transducing specific sgRNAs in A549-Cas9 cells. Clonal cell lines were established post-G418 selection by limiting dilution. Since CMAS is the rate-limiting enzyme in sialic acid biosynthesis, we tested for loss of CMAS function by staining cells with *Sambucus nigra* lectin (SNA), a lectin that specifically binds to α2,6-linked sialic acid on the cell surface, and performing flow cytometry. Five to ten clonal lines were initially screened for loss of SNA staining. The clonal line with the lowest amount of cell surface sialic acid was selected for further validation and infection experiments. Reduction in CMAS protein and *CMAS* mRNA were confirmed by western blot and RT-qPCR (**Figure 1A,B**). SNA binding was abundant on parent A549 cells and markedly decreased in CMAS-deficient cells. Staining with the α2,3-linked sialic acid-binding *Maackia amurensis* lectin (MAA) is less abundant than with SNA in parent A549 cells yet is also reduced in CMAS-deficient cells (**Figure 1C**).

**Figure 1.**
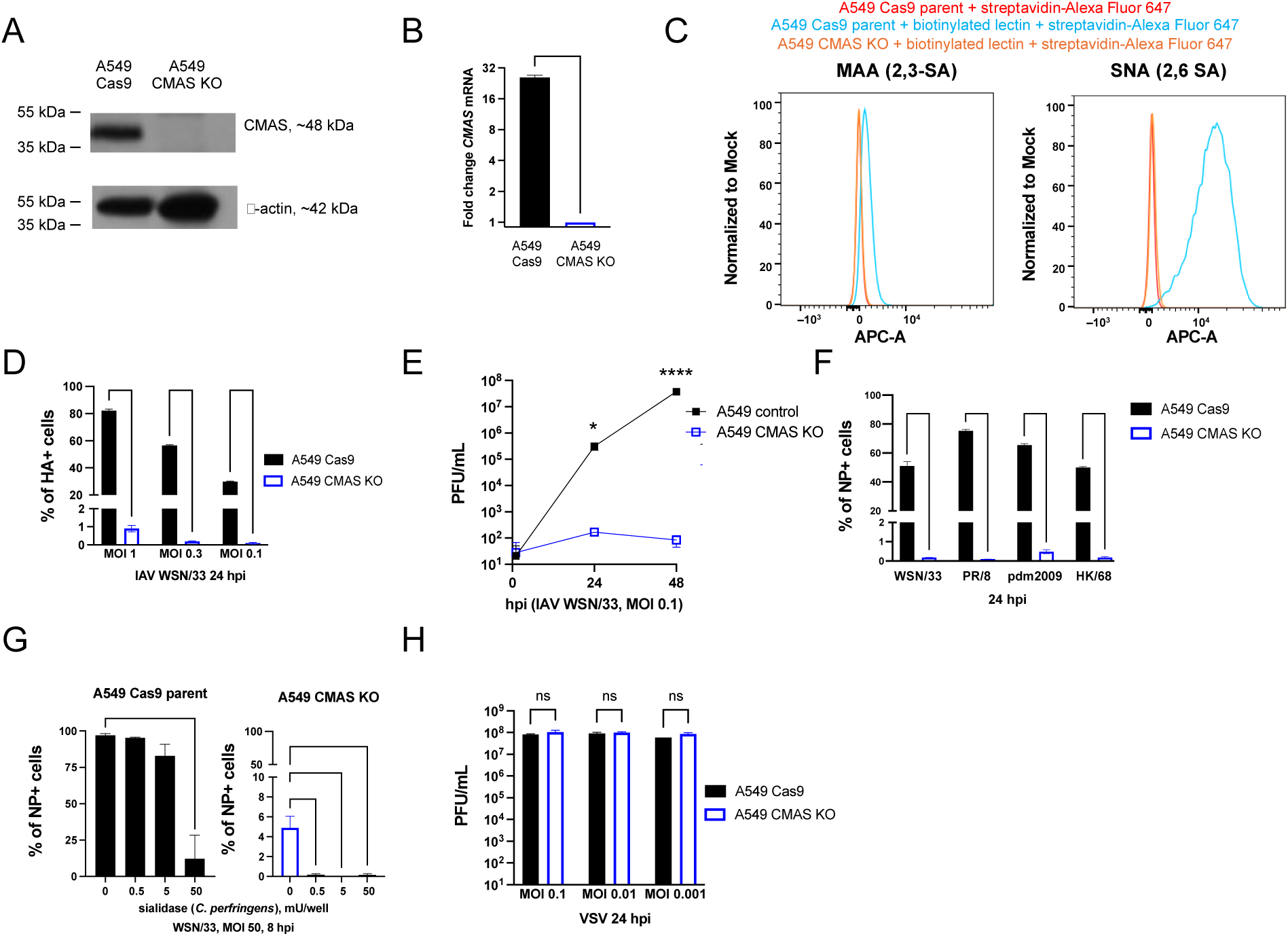
IAV infection is reduced while VSV infection is sustained in A549 CMAS-deficient cells. **A.** Western blot from clonal A549 CMAS-deficient cells show the loss of CMAS protein. Actin is included as a loading control. **B.** *CMAS* expression in A549 CMAS KO cells is reduced compared to parent A549 Cas9 cells by RT-qPCR. RNA was extracted from each cell line and reverse transcribed. *CMAS* and *HPRT* (housekeeping gene) were amplified and the *CMAS* fold change was calculated based on the 2^ΔΔCT^ method. ****, p < 0.0001, unpaired t-test. **C.** Flow cytometry histograms of binding of biotinylated SNA lectin (specific to α2,6-linked sialic acid), biotinylated MAA lectin (specific to α2,3-linked sialic acid), and streptavidin-Alexa Fluor 647 to A549 Cas9 and A549 CMAS KO cell lines. Each group has triplicate samples and a representative sample from each group is shown. ****, p < 0.0001. **D.** A549 Cas9 parent and A549 CMAS KO cells were infected with IAV WSN/33 with the indicated MOI. At 24 hpi, the percentage of HA+ cells was determined by anti-HA+ staining normalized to DAPI staining. ****, p < 0.0001, two-way ANOVA. **E.** A549 and A549 CMAS KO cells were infected with IAV WSN/33 at the indicated MOI. Supernatants were collected at 1, 24, and 48 hpi, and plaque-forming units (PFU) were determined. *, p <0.01, ****; p < 0.0001, unpaired t-test. **F.** A549 Cas9 parent and A549 CMAS KO cells were infected with IAV WSN/33 (H1N1), PR/8 (H1N1), pdm2009 (H1N1), and HK/68 (H3N2) using an inoculum at which ≥40% of A549 Cas9 cells are NP+ at 24 hpi (normalized to DAPI). ****, p < 0.0001, two-way ANOVA. **G.** IAV infection is decreased in both parent and CMAS KO A549 cells following sialidase treatment. Cells were treated with up to 50 mU sialidase per well for 1 h, washed, and then infected with IAV WSN/33 at MOI 50. At 8 hpi, the percentage of NP+ cells was determined. ****, p < 0.0001, one-way ANOVA. **H.** A549 Cas9 parent and A549 CMAS KO cells were infected with VSV at the indicated MOI. Supernatants were collected at 24 hpi and the PFU/mL was determined. ns = not significant, two-way ANOVA. Error bars represent the S.D. for triplicate samples. Each experiment was performed at least two times with consistent results.

To test how CMAS deficiency impacts IAV infection, A549-Cas9 and CMAS KO cells were challenged with IAV WSN/33 and then fixed and stained with an anti-hemagglutinin (HA) antibody at 24 hpi. IAV WSN/33 infection was significantly reduced in CMAS KO compared to parent cells (**Figure 1D**). To confirm that multi-cycle viral replication was also impacted, infected cells were monitored over 48 h with supernatant collected at 1, 24, and 48 hpi (**Figure 1E**). CMAS KO reduced infection of several IAV strains, including PR/8 (H1N1), pdm2009 (H1N1), and IAV strain HK/68 (H3N2) to similar degrees (**Figure 1F**).

Since the CMAS KO cells showed evidence of residual IAV infection, we aimed to determine if sialidase treatment further reduces IAV infection. CMAS KO A549 cells were treated with several doses of sialidase prior to infection. While sialidase treatment did not impact cell adhesion or result in cell toxicity, it resulted in a significant reduction in IAV entry based on NP staining at 8 hpi in both parent and CMAS KO A549 cells (**Figure 1G**).

Finally, to determine if the inhibition was specific for IAV, CMAS KO and parent A549 cells were challenged with vesicular stomatitis virus (VSV) at the indicated MOI. Virus in supernatants collected at 24 hpi was titered by plaque assay on BHK-21 cells. CMAS KO did not significantly impact VSV infection, as comparable levels of virus were recovered from supernatants (**Figure 1H**).

### Overexpression of the previously identified CRISPRa candidate B4GALNT2 inhibits infection with IAV but not VSV

We were interested in following up on previously reported studies from independent CRISPRa screens for host factors performed with other IAV strains. Heaton *et al*. reported that *B4GALNT2* overexpression preferentially impacts α2,3-sialic acid binding by adding N-acetylgalactosamine onto α2,3-sialic acid only and resulted in profound effects on infection with IAV strain PR/8 in A549 cells (29). In contrast, Wong et al. reported that MDCK cells overexpressing B4GALNT2 were equally permissive to IAV WSN/33 compared to parent MDCK cells (49). Therefore, we transduced A549-dCas9-VP64 cells with lentivirus expressing B4GALNT2-specific sgRNA and established a clonal B4GALNT2-overexpressing cell line (CalA^B4GALNT2^) by limiting dilution. We confirmed overexpression of *B4GALNT2* by RT-qPCR and by RNA-sequencing (RNA-seq) (**Figure 2A,B**). Differential expression analysis revealed 15 additional transcripts with significantly increased expression in these cells (**Table S2**).

**Figure 2.**
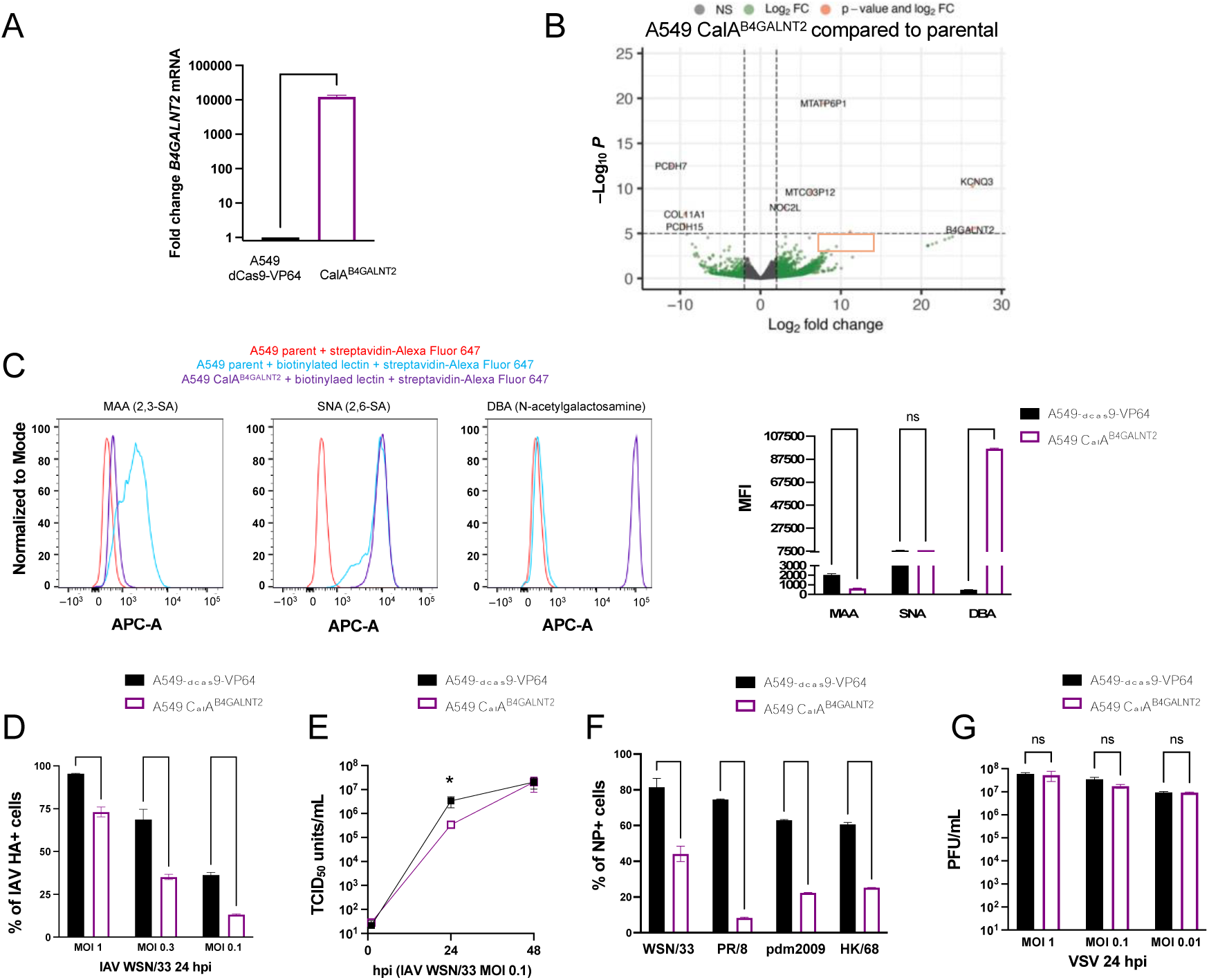
IAV infection is reduced while VSV infection is sustained in A549 CalA^B4GALNT2^ cells. **A.** Expression of *B4GALNT2* is increased in clonal A549 CalA^B4GALNT2^ cells compared to A549-dCas9-VP64 parent cells by RT-qPCR. RNA was extracted from each cell line and reverse transcribed. *B4GALNT2* and *HPRT* (housekeeping gene) were amplified and the *B4GALNT2* fold change was calculated based on the 2^ΔΔCT^ method. ***, p < 0.001, unpaired t-test. **B.** *B4GALNT2* expression is increased with few off-target effects in A549 CalA^B4GALNT2^ cells based on RNA-sequencing and differential expression analysis. Green dots indicate transcripts with log_2_FC >2 or <-2 and -log_10_*P* > 5. **C.** Flow cytometry histograms of binding of biotinylated MAA lectin (specific to α2,3-linked sialic acid), biotinylated SNA lectin (specific to α2,6-linked sialic acid), biotinylated DBA lectin (specific to N-acetylgalactosamine), and streptavidin-Alexa Fluor 647 to A549-dCas9-VP64 and A549 CalA^B4GALNT2^ cell lines. Each group has triplicate samples and a representative sample from each group is shown. Quantification of mean fluorescence intensity (MFI) is shown. ****, p < 0.0001; ns = not significant, one-way ANOVA. **D.** A549-dCas9-VP64 parent and CalA^B4GALNT2^ cells were infected with IAV WSN/33 at the indicated MOI. At 24 hpi, the percentage of HA+ cells was determined by anti-HA staining normalized to DAPI staining. ***, p < 0.001; ****, p < 0.0001, two-way ANOVA. **E.** A549-dCas9-VP64 parent and CalA^B4GALNT2^ cells were infected with IAV WSN/33 at the indicated MOI. Supernatants were collected at 1, 24, and 48 hpi, and viral titers were measured by TCID_50_. *, p < 0.05, unpaired t-test. **F.** A549-dCas9-VP64 parent and CalA^B4GALNT2^ cells were infected with IAV WSN/33, PR/8, pdm2009 (H1N1), and HK/68 (H3N2) using an inoculum at which ≥40% of A549-dCas9-VP64 cells are NP+ at 24 hpi (normalized to DAPI). ****, p < 0.0001; ***, p < 0.001, two-way ANOVA. **G.** A549-dCas9-VP64 and CalA^B4GALNT2^ cells were infected with VSV at the indicated MOI. Supernatants were collected at 24 hpi and PFU/mL was determined. ns = not significant, two-way ANOVA. Error bars represent the S.D. for triplicate samples. Each experiment was performed at least two times with consistent results.

We tested for the effect of B4GALNT2-overexpression on sialic acid by performing lectin staining and flow cytometry as above. As expected, A549 CalA^B4GALNT2^ cells had less MAA (α2,3-sialic acid-specific) binding, no changes in SNA (α2,6-sialic acid-specific) binding, and increased binding to the N-acetylgalactosamine binding lectin *Dolichos biflorus* agglutinin (DBA) compared to parent cells (**Figure 2C**).

Next, A549 CalA^B4GALNT2^ cells were challenged with IAV WSN/33 (H1N1) at the indicated MOI and fixed and stained with an anti-HA antibody at 24 hpi. IAV WSN/33 infection was modestly by significantly reduced in CalA^B4GALNT2^ cells compared to parent A549 cells by HA staining at 24 hpi and by growth assays performed a 48 hpi (**Figure 2D,E**). The cells were then challenged with IAV strains PR/8 (H1N1), pdm2009 (H1N1), and HK/68 (H3N2) (**Figure 2F**) to determine if there was a strain-specific effect. The greatest impairment in IAV infection was seen with the PR/8 strain passaged in embryonated chicken eggs, which is consistent with the findings by Heaton *et al.* (29). The loss of infectivity observed with CMAS deficiency was specific to IAV, as the titers of VSV recovered from the supernatant were similar between these cells and parent cells at 24 h following challenge with VSV at the indicated MOI (**Figure 2G**).

### Loss of CMAS or overexpression of B4GALNT2 prevents binding and entry of IAV

We assessed whether the decrease in IAV infectivity of CMAS KO A549 cells is due to a lack of virus binding. We incubated prechilled parent and CMAS-deficient A549 cells with IAV WSN/33 on ice to facilitate virus binding but not entry. We observed a significant decrease in IAV binding at multiple MOIs tested in CMAS KO cells compared to parent A549 cells. Entry was also restricted, as the fluorescence intensity of IAV WSN/33 vRNA post-entry was significantly reduced in CMAS-deficient cells compared to parent A549 cells as assayed by RNA FISH. Similarly, we tested for IAV WSN/33 binding and entry in A549 CalA^B4GALNT2^ cells by performing virus binding assays. Overexpression of B4GALNT2 resulted in interference with IAV WSN/33 infection through virus binding and internalization, although to a lesser degree than CMAS-deficient cells. Together, these results confirmed that altered expression of these host factors involved in sialic acid biosynthesis restricts IAV infection at the binding and entry stage.

### CRISPRa-mediated overexpression of ADAR1 p150 decreases IAV infectivity

ADAR1 was one of the top ten hits in our CRISPRa screen (**Table S1**). ADAR1 is an enzyme that plays an important role in post-transcriptional editing of RNA. ADAR1 binds to endogenous double-stranded (ds) RNA and catalyzes the hydrolytic deamination of adenosine to inosine. This prevents excessive innate immune responses to dsRNA mediated by the RIG-I-like receptor (RLR) melanoma differentiation-associated protein 5 (MDA5), specifically type I IFN production and ISG as reviewed by Song *et al.* (50). Missense or frameshift mutations in ADAR1 lead to fatal neurodegenerative disorders, such as Aicardi-Goutières syndrome, which is characterized by elevated ISGs due to the failure of the host to discriminate self from non-self dsRNA (51). The role of ADAR1, particularly the IFN-induced p150 isoform, during IAV infection has been examined previously. Ward *et al.* initially reported that the ADAR1 p150 isoform protected host cells from IAV-induced cytopathic effects (52). In a later study, Vogel *et al.* showed that deletion of ADAR1 p150 resulted in sustained RLR signaling and increased apoptosis (53). Of the host factors identified by CRISPRa screening (see **Table S1**), ADAR1 was of interest given that both pro-viral and anti-viral roles have been ascribed to this factor and the consequences of overexpressing the ADAR1 p150 isoform during IAV infection have not been explored. Because the sgRNAs in the library were designed to target the p150 promoter, we elected to overexpress ADAR1 in A549 cells using gRNA specific for the p150 promoter to define its impact on infection by IAV and other viruses.

We transduced three different sgRNAs from Calabrese Library A (the same library used in the initial screen) targeting the ADAR1 p150 promoter into A549-dCas9-VP64 cells (see Materials and Methods) and established clonal cell lines (CalA^ADAR1^) post-selection by limiting dilution. By western blot, we showed that ADAR1 p150 was overexpressed in clonal CalA^ADAR1^ cell lines with guide 1 (clones 7 and 4) and guide 2 (clones 1 and 10). Overexpression of the p110 isoform is observed in all CalA^ADAR1^ clonal cell lines by western blot, which may result from leaky ribosome scanning during transcription (54). Upregulation of *ADAR1* was observed in all clones by RT-qPCR using non-isoform-specific primers. We also detected upregulation of *ADAR1* by next-generation sequencing of CalA^ADAR1^ ^g1^ ^cl7^ cells. Off-target effects in these cells were more prevalent than for *B4GALNT2* overexpressing cells (**Figure 4C**, **Table S3**). Given the potential for off-target effects, we examined three different A549 CalA^ADAR1^ clones (A549 CalA^ADAR1^ ^g1^ ^cl7^, A549 CalA^ADAR1^ ^g2^ ^cl1^, and A549 CalA^ADAR1^ ^g3^ ^cl3^) in our IAV infection studies.

**Figure 3.**
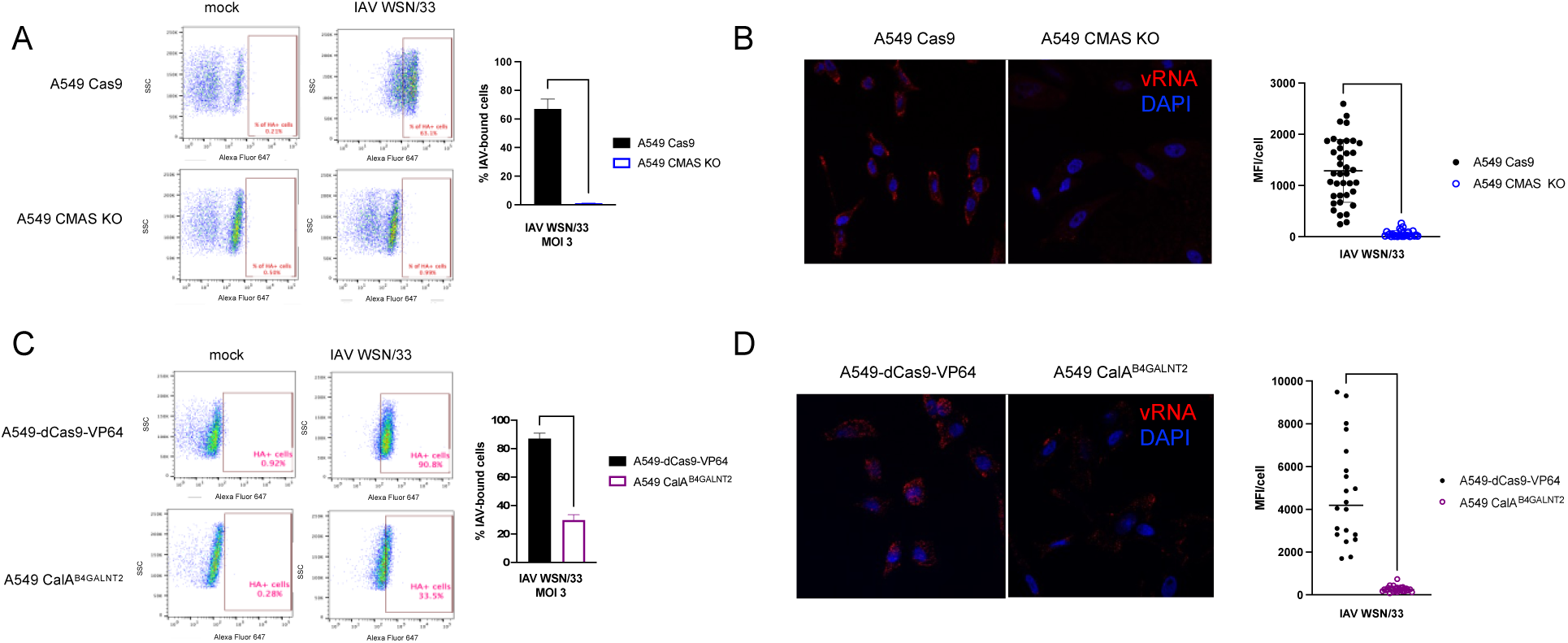
IAV binding and entry are decreased in A549 CMAS KO cells and in A549 CalA^B4GALNT2^ cells compared to parent A549 cells. **A.** Representative flow cytometry plots with quantification of IAV-bound A549 Cas9 parent and A549 CMAS KO cells. Cells were challenged with IAV WSN/33 at the indicated MOI (or mock-infected) for 1 h at 4 °C to facilitate binding but not allow for endocytosis. Cells were washed, fixed, and then stained with anti-HA monoclonal antibody followed by Alexa Fluor 647-conjugated secondary antibody. IAV-bound cells were identified based on comparison to anti-HA staining of mock-infected cells. All data were analyzed with FlowJo software. Quantification of the percentage of IAV-bound A549 Cas9 parent and A549 CMAS KO cells at the indicated MOI is shown. Error bars represent the S.D. for triplicate samples. ***, p < 0.001, two-way ANOVA. **B.** Representative composite immunofluorescence images taken at 630x magnification for A549 Cas9 parent and A549 CMAS-deficient cells challenged with IAV WSN/33 at MOI 3 and fixed 20 min post-infection. IAV entry was defined by WSN/33 segment 7 vRNA staining by FISH (red) and cell nuclei were defined by DAPI staining (blue). The average mean fluorescence intensity (MFI, representing the average intensity of red staining) per cell was quantified and is shown in the graph. Each dot represents one cell and the horizontal bar shows the mean value. Three images with at least eight cells/field of view were captured for each cell line. ****, p < 0.001, Mann-Whitney test. **C.** Representative flow cytometry plots with quantification of IAV WSN/33-infected A549-dCas9-VP64 parent and A549 CalA^B4GALNT2^ cells. The approach was similar to that shown in A. **D.** Representative composite immunofluorescence images taken at 630x magnification for A549-dCas9-VP64 parent and A549 CalA^B4GALNT2^ cells. The approach was similar to that for B.

**Figure 4.**
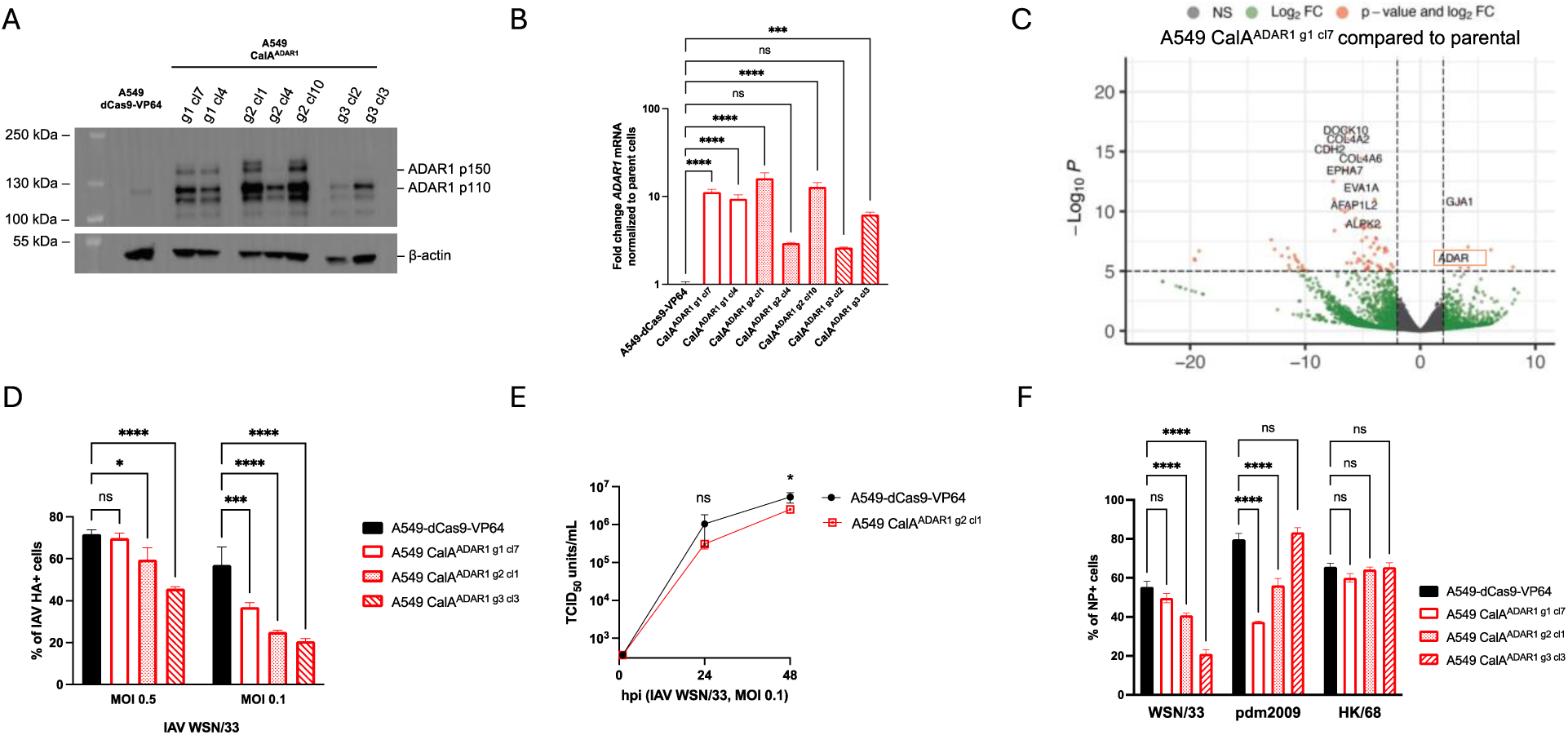
ADAR1 overexpression in A549 CalA^ADAR1^ clonal cells is associated with a modest reduction in IAV infection. **A.** Equal protein concentrations of CalA^ADAR1^ clonal cells and A549-dCas9-VP64 parent cells were loaded onto a 4-8% Tris-Acetate gel for SDS-PAGE electrophoresis and transferred to a PVDF membrane, which was then probed with antibodies against ADAR1 and actin. Chemiluminescent substrate was added and the blot was imaged. **B.** Expression of *ADAR1* expression in clonal A549 CalA^ADAR1^ cells is increased compared to A549-dCas9-VP64 parent cells by RT-qPCR. RNA was extracted from each cell line and reverse transcribed. *ADAR1* and *HPRT* (housekeeping gene) were amplified and the *ADAR1* fold change was calculated based on the 2^ΔΔCT^ method. The experiment was performed twice. ***, p < 0.001; ****, p < 0.0001; ns = not significant, one-way ANOVA. Error bars represent the S.D. for triplicate samples. The primers for ADAR1 bind to exon 2 which is expressed in both p110 and p150 isoforms. **C.** *ADAR1* expression is increased with some off-target effects in A549 CalA^ADAR1^ ^g1^ ^cl7^ cells based on RNA-sequencing and differential expression analysis. Transcripts with log_2_FC >2 or <-2 are green and those with -log_10_*P* > 5 are orange. **D.** A549-dCas9-VP64 parent and A549 CalA^ADAR1^ clonal cells were infected with IAV WSN/33 at the indicated MOI. At 24 hpi, the percentage of HA+ cells was determined by anti-HA staining normalized to DAPI staining. *, p < 0.05; ***, p < 0.001; ****, p < 0.0001; ns = not significant by two-way ANOVA. **E.** A549-dCas9-VP64 parent and A549 CalA^ADAR1^ ^g2^ ^cl1^ cells were infected with IAV WSN/33 at the indicated MOI. Supernatants were collected at 1, 24, and 48 hpi and viral titers were by TCID_50_. *, p < 0.05; ns = not significant by unpaired t test. **F.** A549-dCas9-VP64 parent and CalA^ADAR1^ cells were infected with IAV WSN/33, pdm2009 (H1N1), and HK/68 (H3N2) using an inoculum at which ≥40% of A549-dCas9-VP64 cells are NP+ at 24 hpi (normalized to DAPI). ***, p < 0.001; ****, p < 0.0001; ns = not significant, two-way ANOVA. Error bars represent the S.D. for triplicate samples. Each experiment was performed at least two times with consistent results.

Three CalA^ADAR1^ clonal cell lines and parent A549-dCas9-VP64 cells were challenged with IAV WSN/33 at two different MOI, and then 24 hours later were fixed and stained for HA to determine the percentage of infection. Interestingly, IAV WSN/33 was significantly but only modestly inhibited as determined by HA staining of these various A549 CalA^ADAR1^ cell lines (**Figure 4D**). Moreover, a viral growth assay quantifying the IAV WSN/33 TCID_50_ over 48 h showed a reduction in titers of less than 10-fold for A549 CalA^ADAR1^ ^g2^ ^cl1^ cells compared to parent cells (**Figure 4E**) and trends with IAV pdm2009 and HK/68 strains were inconsistent with those for WSN/33 (**Figure 4F**). Altogether, these data do not support a strong anti-IAV role for ADAR1, either p110 or p150, in A549 cells.

### Overexpression of ADAR1 results in increased infection of CVB but not VSV

To determine if ADAR1 overexpression impacts other viruses, the A549 CalA^ADAR1^ ^g1^ ^cl7^ clonal cell line was challenged with VSV at the indicated MOI for 24 h. Supernatants were collected and titered on BHK-21 cells. No difference in the VSV titers was observed from the supernatant of CalA^ADAR1^ cells compared to that from the parent A549-dCas9-VP64 cells (**Figure 5A, left**). On the other hand, ADAR1 overexpression was robustly pro-viral during CVB4 infection, as a 10-fold increase in viral titers was observed in the supernatant from two different CalA^ADAR1^ cell lines compared to A549-dCas9-VP64 cells at 24 hpi (**Figure 5A, right**). To test how ADAR1 deficiency impacts CVB replication, we challenged HEK293T WT and HEK293T ADAR1 KO cells lacking both the p110 and p150 isoforms (31) with CVB4 strain JVB. As expected, the absence of ADAR1 resulted in a significant decrease in CVB4 titers at 24 hpi but had no effect on VSV replication (**Figure 5B**). Thus, our data support a pro-viral role for ADAR1 during CVB4 infection.

**Figure 5.**
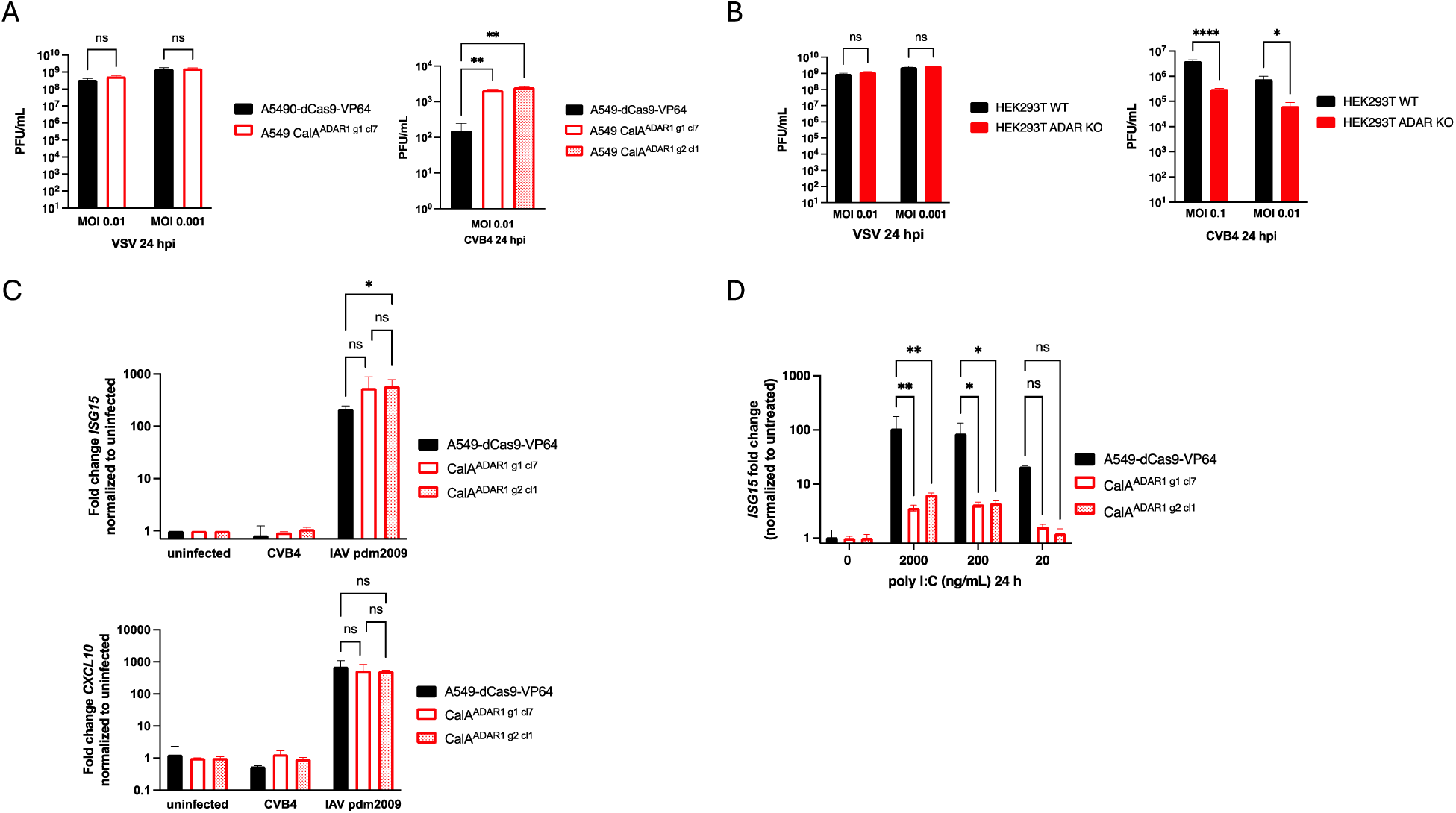
CVB4 replication is increased in A549 CalA^ADAR1^ clonal cells and the interferon-stimulated gene response is absent in response to CVB infection in both A549 parent and A549 CalA^ADAR1^ clonal cells. **A.** A549-dCas9-V64 parent and A549 CalA^ADAR1^ cells were infected with VSV or CVB4 at the indicated MOI. At 24 hpi, supernatants were collected and titered. ns = not significant by one-way ANOVA for VSV. **, p < 0.01 by one-way ANOVA for CVB4. **B.** HEK293T parent and ADAR1 KO cells were infected with VSV or CVB4 at the indicated MOI. At 24 hpi, supernatants were collected and titered. Error bars represent the S.D. for triplicate samples. ns = not significant for VSV. ****, p < 0.0001; *, p <0.05 by one-way ANOVA for CVB4. **C.** A549-dCas9-VP64 parent and A549 CalA^ADAR1^ cells were infected with CVB4 at MOI 0.1 or IAV pdm2009 at MOI 0.3 for 24 h. RNA was extracted from each cell line and reverse transcribed. *ISG15, CXCL10,* and *HPRT* (housekeeping gene) were amplified, and the *ISG15 or CXCL10* fold change was calculated based on the 2^ΔΔCT^ method. *, p < 0.05; ns = not significant, one-way ANOVA. Error bars represent the S.D. for triplicate samples. **D.** A549-dCas9-VP64 parent and A549 CalA^ADAR1^ cells were treated with dsRNA mimetic poly I:C at the indicated concentrations for 24 h. *ISG15* and *HPRT* (housekeeping gene) were amplified and the *ISG15* fold change was calculated based on the 2^ΔΔCT^ method. **, p < 0.01; *, p < 0.05; ns = not significant, one-way ANOVA. Error bars represent the S.D. for triplicate samples. Each experiment was performed at least two times with consistent results.

Next, we considered whether the pro-viral effect of *ADAR1* overexpression during CVB4 infection is linked to a diminished IFN response. Because IFN production following CVB infection is largely mediated by MDA5 (55), we hypothesized that ADAR1 overexpression diverts dsRNA replicative intermediates from engaging with MDA5. Thus, we challenged A549-dCas9-VP64 parent cells and A549 CalA^ADAR1^ cells with CVB4 and extracted RNA from the cells at 24 hpi to evaluate ISG expression using RT-qPCR. Unexpectedly, *ISG15* and *CXCL10* are not induced following CVB4 infection of either CalA^ADAR1^ or A549-dCas9-VP64 cells (**Figure 5C**); this result was robust and consistent but surprising given that CVB4 induces ISGs in other cell systems such as human islets (56, 57). To exclude a generalized defect in ISG production, A549-dCas9-VP64 parent and CalA^ADAR1^ cells were infected with IAV pdm2009 and RNA was extracted at 24 hpi for RT-qPCR. *ISG15* and *CXCL10* are highly and equally induced in A549 CalA^ADAR1^ and A549-dCas9-VP64 cells (**Figure 5C**), confirming that the ISG response in A549 CalA^ADAR1^ cells is intact. We also treated cells with the dsRNA mimetic poly I:C to induce an ISG response. At 24 h post-treatment, *ISG15* induction is present but significantly dampened in CalA^ADAR1^ cells compared to A549-dCas9-VP64 cells (**Figure 5D**). Here, poly I:C may be sequestered by ADAR1 and diverted away from detection by MDA5. Further studies in diverse host cells are needed to determine whether such findings are specific to A549 cells or can be observed more broadly.

## Discussion

In this study, we demonstrate that the roles of host genes involved in productive virus infection can be identified either through either knockout, as exemplified by *CMAS*, or overexpression, as demonstrated by *B4GALNT2* and *ADAR1*. CMAS is an essential enzyme in the sialic acid biosynthesis pathway catalyzing the rate-limiting step of converting N-acetylneuraminic acid to 5-monophosphate N-acetylneuraminic acid in the nucleus, which is an essential building block for sialic acid (45). CRISPR-Cas9-mediated deletion of *CMAS* disrupts sialic acid biosynthesis, resulting in minimal production of α2,3- and α2,6-linked sialic acids on the cell surface. Consequently, we demonstrate that CMAS KO cells are less permissive to IAV infection due to impaired virus binding, and that sialidase treatment eliminates any residual infection.

*B4GALNT2* is responsible for the addition of N-acetylgalactosamine onto the subterminal galactose of the α2,3-linked sialic acid monosaccharide. *B4GALNT2* was a positive hit in the CRISPRa screen as previously reported (29), despite the use of different IAV strains in this study. We observed varying levels of resistance to infection by different IAV strains in CalA^B4GALNT2^ cells, perhaps due to the preference of each strain for either α2,3- or α2,6-linked sialic acid. Since overexpression of *B4GALNT2* specifically modifies α2,3-linked sialic acid, CalA^B4GALNT2^ cells can be used to distinguish the dependency of different viruses to either α2,3- or α2,6-linked sialic acids. In future studies, it would be interesting to confirm whether overexpression of *B4GALNT4,* a gene within the galactosaminyltransferase family, is involved in IAV infection. While B4GALNT2 transfers N-acetylgalactosamine in a β1,4-linkage from UDP-N-acetylgalactosamine to the sub-terminal galactose of α2,3-sialyllactosamine, B4GALNT4 instead catalyzes the addition of N-acetylgalactosamine to underlying N-acetylglucosamine. B4GALNT4 overexpression may negatively impact IAV WSN/33 infection since it was identified in our CRISPRa screen yet has been reported as a potential pro-viral factor in a knockout screen (13).

ADAR1 catalyzes the deamination of adenosine to inosine (A-to-I) on dsRNA, thereby preventing aberrant IFN secretion in the absence of virus infection by masking endogenous dsRNA from detection by RLRs. ADAR1 was identified in our CRISPRa screen as possibly anti-viral when overexpressed. ADAR1 can function as either a pro- or anti-viral host factor depending on the virus (48). In the context of IAV infection, ADAR1 p150 has been reported to be a pro-viral host factor functioning to block RLR signaling and apoptosis, and ADAR1 p110 has been shown to be an anti-viral host factor (53). ADAR1 can also act as an anti-viral host factor through A-to-I editing of matrix (M) mRNA (58). However, specific overexpression of ADAR1 in A549 cells had very little inhibitory effect during IAV WSN/33 infection. Cells with guide 3, which appeared to increase p110 but not p150 by western blot, had a greater reduction of IAV WSN/33 infection compared to those with guide 1 or 2. This is consistent with the report by Vogel et al. that ADAR1 p110 expression hinders IAV replication and may be responsible for the original phenotype observed with ADAR1-overexpressing cells in the CRISPRa screen (53).

Interestingly, we found that ADAR1 overexpression had a pro-viral effect during infection with CVB4. Two prototypical ISGs, *ISG15* and *CXCL10,* are not induced in parent or ADAR1-overexpressing A549 cells following CVB4 challenge. IFN signaling and ISG induction are intact in both parent and ADAR1-overexpressing cells, as *ISG15* and *CXCL10* are induced robustly and equally following IAV challenge. However, reduced expression of ISGs in poly I:C-challenged ADAR1-overexpressing A549 cells may be due to ADAR1 diverting poly I:C away from downstream interactions with MDA5, but this did not appear to be the case for CVB4. Pro-viral effects of ADAR1 overexpression during CVB4 infection could be due to ADAR1 interactions with protein kinase R (PKR), as ADAR1 p150 suppresses PKR activation by competitive binding to dsRNA (51, 52), while a functional PKR is a classical cytosolic dsRNA sensor that shuts off viral protein synthesis and reduces virus replication (59). Since both ADAR1 p150 and ADAR1 p110 are induced in our CRISPRa system, despite our design to specifically upregulate the p150 isoform, future studies should express these isoforms from cDNAs to decouple the contribution of each in the context of CVB4 and IAV infection. Additionally, blockade of PKR could be used to confirm its possible role in ADAR1-driven pro-CVB4 effects.

In sum, our work demonstrates that CRISPR-based modulation strategies can identify host factors that could serve as novel targets for antiviral strategies against IAV. The assays we developed help elucidate the role of such host factors during the infection process. Further, they provide insight into how these factors contribute to infection with other viruses.

## Acknowledgments

This work is dedicated to the late Dr. Robert Finberg.

## Funding

This work was supported by the Defense Advanced Research Projects Agency, contracted via the Department of Navy, Office of Naval Research under the Federal Award Number N660011924036 (W.M.M., K.A.F, J.P.W.).

## Authors’ contributions

P.P.K., W.M.M., N.S., K.A.F., and J.P.W. conceived the study. P.P.K., P.L, Z.Z.J., E.S.B., T.C., and W.M.M performed the experiments. P.P.K., M.I.T., N.S., and J.P.W. wrote the manuscript. J.P.W. and K.A.F. acquired the funding. J.P.W. is the guarantor.

## Supplementary Material

**Figure S1.**
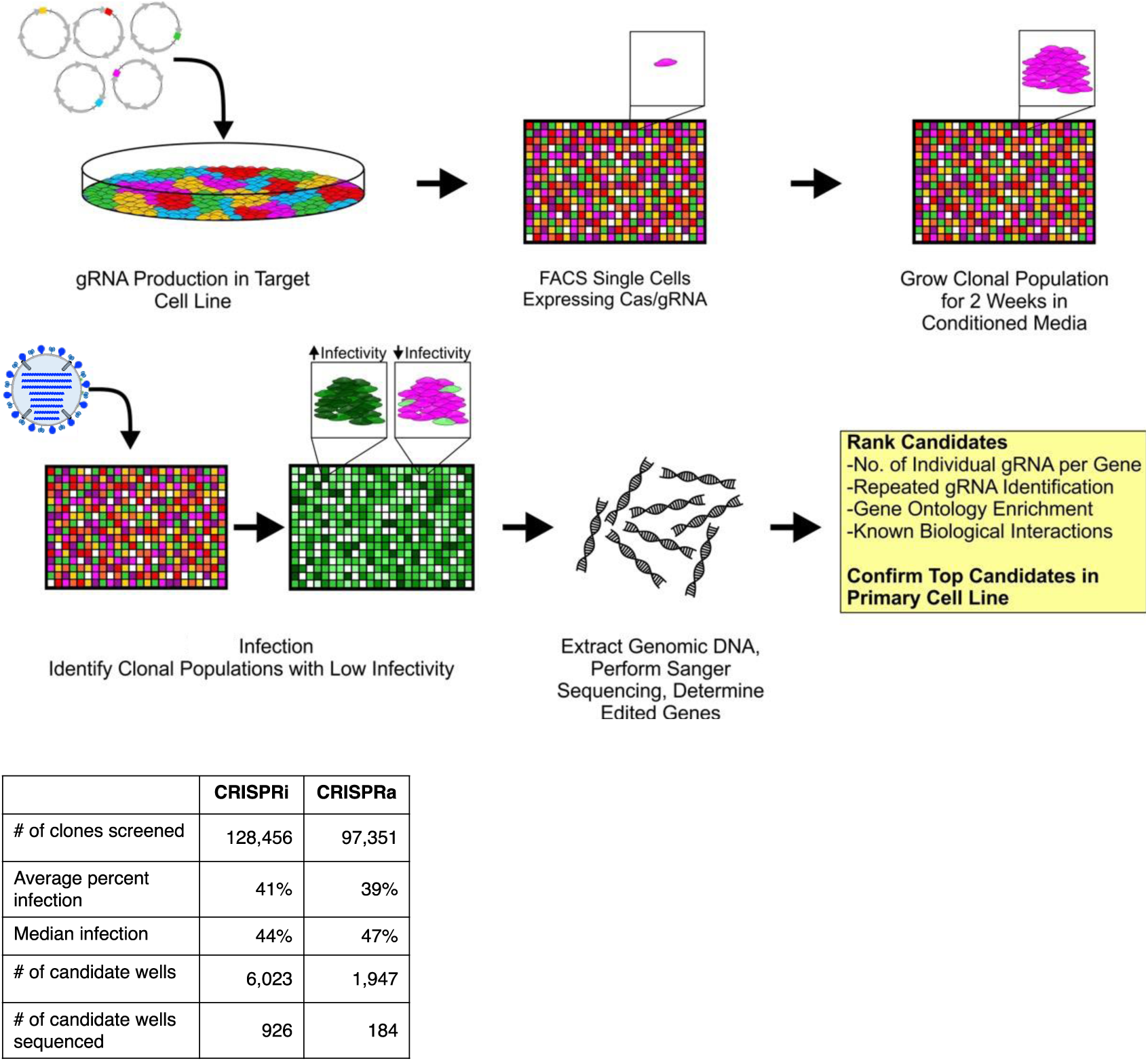
Schematic of arrayed CRISPRa/i screening strategy for host genes that impact IAV infection and the strategy for candidate prioritization and analysis. Candidates from the screen were assessed for common biological pathways and prioritized based on multiple metrics, including repeated gRNA identification, gene ontology enrichment, gene expression databases (e.g., the Genotype-Tissue Expression), known biological interactions, and other publicly available data. The data were also compared to the bulk RNA-Seq data for baseline levels of genes, as well as differentially regulated genes following viral infection.

**Table S1.**
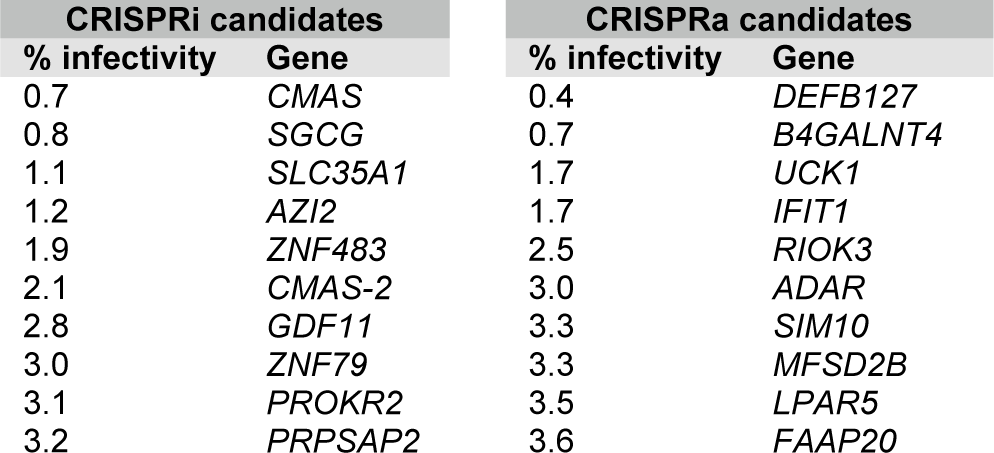
Top ten CRISPRi and CRISPRa candidates yielding decreased infectivity of IAV.

**Table S2.**
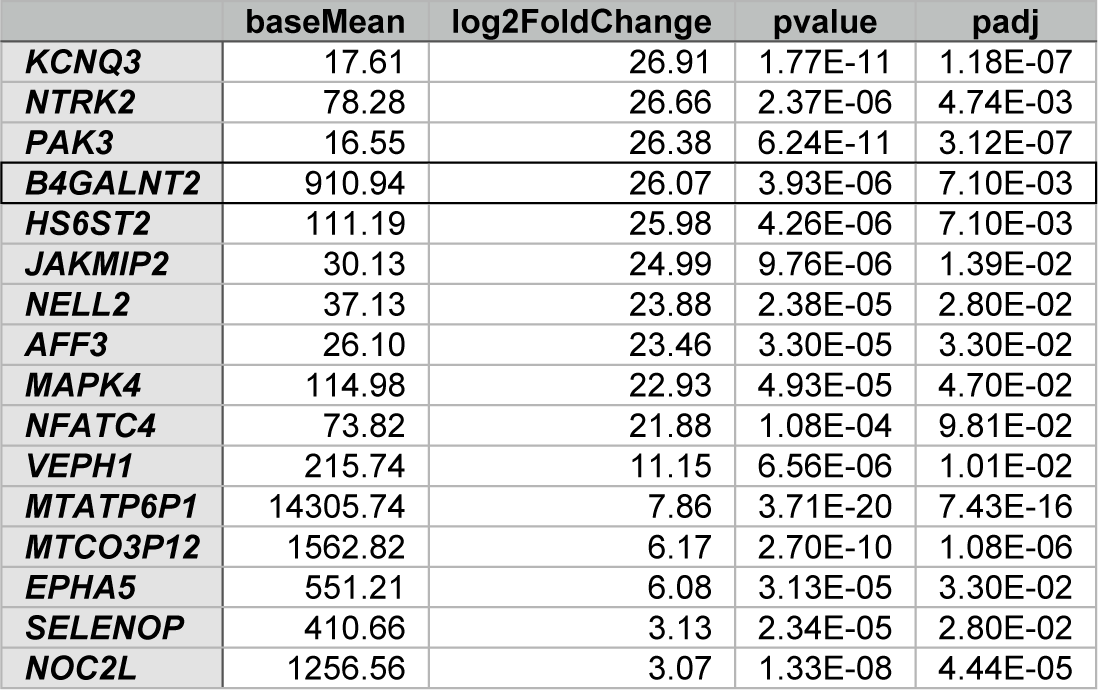
RNA-sequencing differential expression for A549 CalA^B4GALNT2^ cells compared to parent cells, log_2_ fold change >2.5.

**Table S3.** RNA-sequencing differential expression for A549 CalA^ADAR1^ cells compared to parent cells, log_2_ fold change>2.5.

|  | baseMean | log2FoldChange | pvalue | padj |
| --- | --- | --- | --- | --- |
| **ZNF700** | 15.84 | 8.50 | 6.48E-04 | 5.06E-02 |
| **ZNF136** | 16.18 | 8.16 | 3.10E-04 | 2.97E-02 |
| **OTOF** | 15.85 | 8.16 | 3.41E-04 | 3.19E-02 |
| **COL11A1** | 41.63 | 8.06 | 4.57E-06 | 1.07E-03 |
| **ZNF91** | 72.03 | 6.13 | 1.68E-07 | 6.97E-05 |
| **ZSCAN18** | 36.83 | 5.60 | 1.03E-03 | 7.03E-02 |
| **PCDH7** | 100.48 | 5.44 | 5.61E-04 | 4.67E-02 |
| **CHGB** | 8093.17 | 5.28 | 4.65E-05 | 6.59E-03 |
| **CALCRL** | 126.78 | 5.03 | 6.18E-04 | 4.91E-02 |
| **GUCY1A2** | 70.09 | 4.87 | 2.25E-04 | 2.25E-02 |
| **SBK2** | 156.16 | 4.83 | 3.60E-04 | 3.33E-02 |
| **ZNF420** | 33.45 | 4.83 | 4.58E-04 | 4.04E-02 |
| **IGSF10** | 478.06 | 4.65 | 1.64E-03 | 9.87E-02 |
| **SERPINI1** | 760.60 | 4.19 | 5.01E-06 | 1.14E-03 |
| **COL26A1** | 135.37 | 4.16 | 9.57E-08 | 4.90E-05 |
| **PTCH1** | 1146.09 | 3.78 | 1.81E-05 | 3.04E-03 |
| **SNTB1** | 326.84 | 3.54 | 2.99E-05 | 4.57E-03 |
| **BDKRB1** | 87.98 | 3.51 | 1.15E-03 | 7.78E-02 |
| **DLK2** | 102.74 | 3.48 | 5.13E-06 | 1.15E-03 |
| **GJA1** | 8959.22 | 3.43 | 1.52E-11 | 2.35E-08 |
| **PDE3B** | 734.23 | 3.39 | 1.35E-04 | 1.51E-02 |
| **HOXC8** | 172.02 | 3.22 | 4.11E-04 | 3.72E-02 |
| **PCDH1** | 159.87 | 3.17 | 4.95E-04 | 4.23E-02 |
| **PARM1** | 499.23 | 2.97 | 2.23E-05 | 3.65E-03 |
| **CA11** | 549.82 | 2.90 | 5.03E-04 | 4.26E-02 |
| **ADAR** | 107586.14 | 2.88 | 7.84E-07 | 2.61E-04 |
| **HOXB8** | 307.58 | 2.55 | 3.30E-04 | 3.12E-02 |
| **COL12A1** | 1978.50 | 2.53 | 1.27E-03 | 8.34E-02 |
| **RAP1GAP2** | 2318.42 | 2.51 | 8.48E-04 | 6.11E-02 |

